# Natural language-based representation and modeling of RBP binding

**DOI:** 10.64898/2026.02.05.704032

**Authors:** Shaimae I. Elhajjajy, Zhiping Weng

## Abstract

RNA-binding proteins (RBPs) are critical regulators of the human transcriptome, but the binding patterns of most RBPs are insufficiently characterized. While sequence context facilitates RBP binding specificity, its precise contribution remains unclear. Existing computational methods to decipher RBP binding patterns are limited by their architecture-dependence, challenging interpretability, and, importantly, lack of focus on context. We present a novel comprehensive approach to address the aforementioned knowledge gaps. We first introduce a natural language-based representation to model RNA sequences using lexical, syntactic, and semantic forms, then devise a sequence decomposition method based on these structures to deconstruct RNA sequences into regions, each containing a target ***k***-mer and its flanking contexts. We leverage this linguistic conceptualization to predict RBP binding under a Multiple Instance Learning (MIL) framework, which we solve using a novel method of significant region extraction termed “iterative relabeling”. We demonstrate that our bottom-up approach discovers key regions contributing to RBP binding in an architecture-dependent, accurate, and interpretable manner.

## 1 Introduction

RNA-binding proteins (RBPs) dynamically regulate RNA processing to modulate accurate gene expression and sufficient transcriptomic diversity, both of which are essential to cell survival. The majority of the 1,500 to 2,000 human RBPs remain insufficiently characterized in their binding patterns, activities, and functions [1, 2]. While RBPs with structurally diverse RNA-binding domains can recognize short, simple sequence motifs that often bear striking resemblance to one another, they remain capable of binding distinct transcriptomic sites [3]. Previous findings have demonstrated that sequence context facilitates RBP binding specificity [3], but these findings have not been systematically validated at large scale *in vivo*. An improved understanding of the role of sequence context in facilitating specific RBP binding will shed light on RBP binding patterns and functions in RNA regulation.

Current approaches primarily rely on architecture-dependent approaches for sequence-based genomic prediction of transcription factor (TF) and RBP binding. However, these models are notoriously difficult to interpret, resulting in a lack of transparency regarding the features learned and used for decision-making [4]. Due to their architecture-dependence, these models also rely on tracking activation values across layers in order to derive feature importance. Further, the performance-interpretability tradeoff remains a challenge, which can lead to reports of satisfactory model prediction accuracy but erroneous motifs for well-characterized RBPs. Importantly, existing computational approaches focus heavily on RBP binding motif discovery but have rarely investigated the contextual determinants of RBP binding. Hence, there remains a need for an RBP binding prediction approach that simultaneously addresses these technical limitations and enhances our understanding of the underlying mechanisms regulating RBP binding specificity.

Conventional approaches of discovering motifs tend to use methods that treat motif instances in isolation, without considering the contribution of neighboring *k*-mers; because the influence of sequence context is ignored, this could lead to the identification of false motif instances. In contrast, we introduce a context-aware approach that is built around a central, candidate target *k*-mer surrounded by its flanking regions. This formulation allows us to deconstruct sequences into regions, identify the regions that are most significant, validate the candidacy of motif instances within these significant regions, and then concentrically reconstruct sequences in a bottom-up fashion, thus providing greater resolution in the discovery of RBP motifs and their surrounding contexts.

Our linguistic conceptualization led to an RBP binding model under the Multiple Instance Learning (MIL) framework [5]. In this formulation, we treat sequences as “bags” with known labels, and regions as “instances” of bags with unknown labels; our objective is to determine which of the region(s) (i.e., key instance[s]) within a bound sequence significantly contribute to its positive label [6, 7]. Since its inception in medicinal chemistry, MIL has been applied to many diverse fields; this is, to the best of our knowledge, the first formulation of the RBP binding prediction task as an MIL problem.

We present herein a novel method that integrates principles from natural language, linguistics, and weak supervision to model RBP binding patterns. We define a lexical, syntactic, and semantic representation of RNA sequences that clearly distinguishes between RBP motif and context *k*-mers. Our linguistic formulation naturally lends the RBP binding prediction task to be cast under an MIL framework, which we solve with our significant region extraction method termed “iterative relabeling” that is analogous to key phrase extraction methods in natural language processing. We demonstrate that our models yield strong, robust performance across an array of RBPs in both HepG2 and K562. Crucially, our predictions enable the accurate discovery of RBP binding motifs and contexts. Taken together, our work presents a novel, interpretable, and biologically relevant approach for natural language-based modeling of RBP binding that is architecture-dependent, versatile, and generalizable, thus providing a new avenue by which to improve our understanding of RBP-mediated RNA regulation.

## 2 Methods

### 2.1 Problem Definition, Assumptions, Requirements, and Objectives

Given a set of labeled sequences comprising a subset of bound sequences labeled positive and a subset of unbound sequences labeled negative, the problem is to accurately pinpoint RBP binding sites within each positive sequence under the following assumptions:

1. Each positive sequence represents a putative RBP binding site.
2. *k*-mers contributing to RBP binding events are enriched in positive sequences.
3. Enriched *k*-mers within positive sequences contribute to the RBP binding event either (a) as a motif instance recognized and bound by the RBP or (b) as a contextual element within the flanking RNA sequence that facilitates the binding event.
4. Positive sequences contain a greater abundance of enriched *k*-mers relative to negative sequences.
5. Enriched *k*-mers form a region significant to binding.
6. There may be more than one significant region per sequence.

Our objective is to investigate the identity and roles of enriched *k*-mers comprising RBP binding sites and their flanking regions. Accordingly, we required a comprehensive method that enables:

1. A simple representation of sequences into their basic constitutive units (analogous to words as lexical units).
2. Decomposition of sequences into their constitutive regions (analogous to syntactic forms in natural language).
3. Determination of the meaning, significance, contribution, and roles of these constitutive regions to the structure and composition of their respective sequences.
4. Determination of the meaning, significance, contribution, and roles of the basic constitutive units to their respective regions and sequences (analogous to lexical semantics).
5. Reconstruction of sequences from their constitutive units and regions.

Finally, we also required our method to provide high resolution, interpretability, flexibility, easy access to genomic coordinates, and architecture independence.

### 2.2 Dataset composition

We downloaded merged replicate enhanced crosslinking and immunoprecipitation (eCLIP) peak files in HepG2 and K562 from the ENCODE portal [8, 9] (**Supplemental Table S1**). For each RBP, we defined positive training sequences as 101 nucleotide (nt) centered peaks that (1) originated from chromosomes {chr1-22, chrX}, (2) contained only the canonical nucleotides {A, C, G, U}, and (3) are located in any of the genomic regions {tRNA, miRNA, miRNA-proximal, CDS, 3’UTR, 5’UTR, 5’SS, 3’SS, proximal intron, distal intron, noncoding exon}, assigned according to the strategy in [9]. To ensure matching nucleotide content and balanced cross-validation groups, we defined negative sequences by randomly sampling 101 nt sequences from the same genomic regions and chromosomes as our positive sequences, resampling as needed to prevent any overlap with the positive set. We then pooled our positive and negative sequences to form our complete training dataset. We assigned each sequence in our dataset to one of 12 cross-validation groups according to the chromCV method [10]; the largest of these groups was set aside as the held-out independent test set.

### 2.3 A linguistics- and natural language-based representation of RBP binding sequences

DNA and RNA are believed to possess an underlying language, whose structure and components remain elusive [11]. The core components of any natural language comprise a lexicon, syntactic rules, and semantic rules that permit lexical, syntactic, and semantic analysis to understand its meaning (**Figure 1A**). Analogous structures and rules in RNA remain enigmatic, rendering genomic lexicography, syntax, and semantics as nontrivial endeavors. Without a genomic “delimiter” assuming the role of a language delimiter, and without knowledge of permissible word lengths, enumeration of all possible *k*-mers becomes far too sparse and unwieldy due to the exponential nature of combinatorics (**Supplemental Note .1**).

**Fig. 1.**
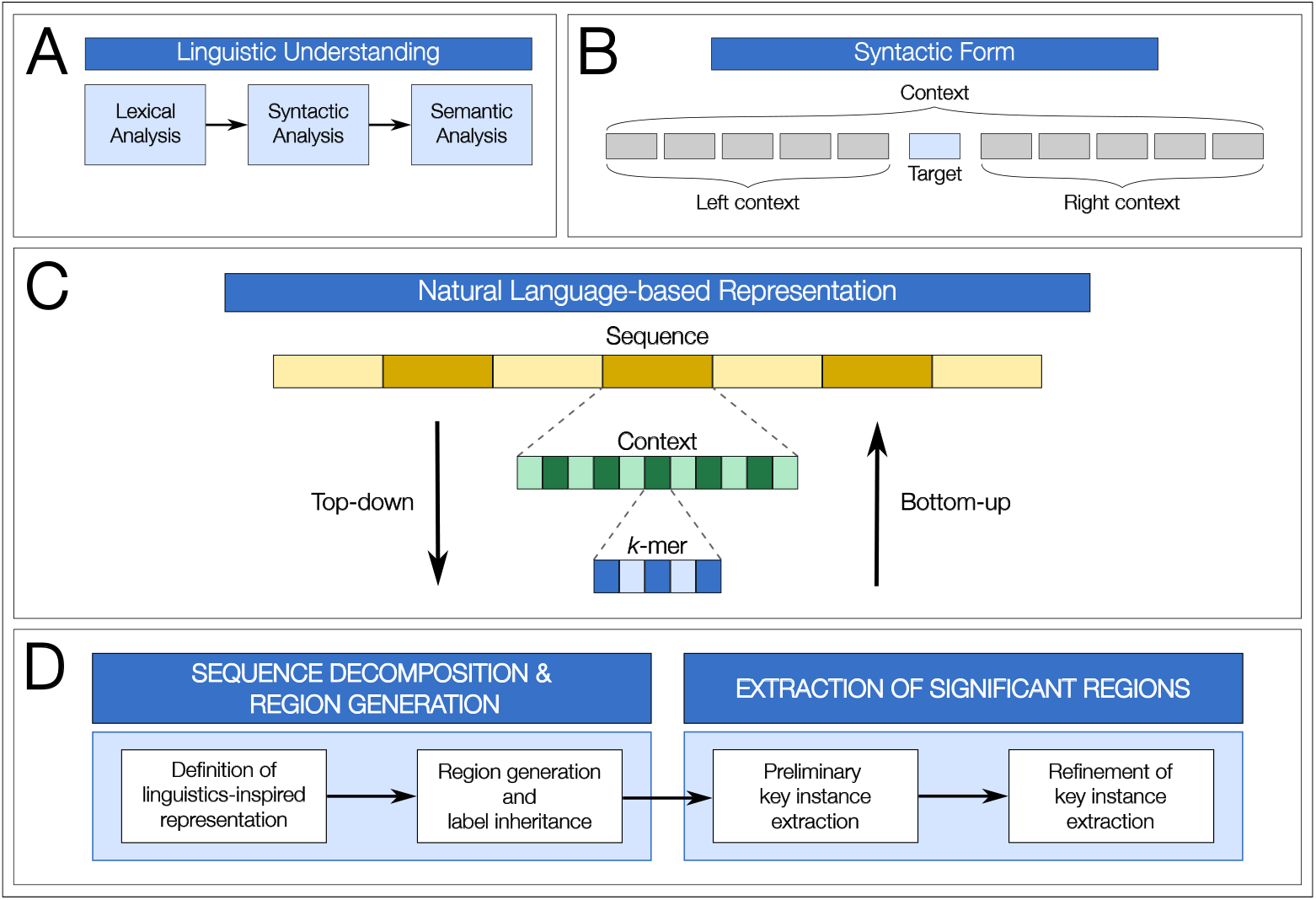
Linguistic-inspired conceptual formulation and structural design. (A) Natural language is understood at different levels of linguistic analysis: lexical, syntactic, and semantic. (B) Our proposed syntactic form consisting of a central “target” *k*-mer flanked by two segments, each containing a pre-determined number of *k*-mers, termed the “left context” and “right context’, respectively. (C) High-level overview of our sequence decomposition strategy, in which sequences are deconstructed into context and *k*-mers. (D) High-level overview of our full framework’s conceptualization. After defining our linguistics-inspired representation, we perform sequence decomposition to generate regions and assign region labels using label inheritance (left). Next, we perform a two-phase significant region extraction, consisting of preliminary and refinement phases, in which we identify key instances (i.e., significant regions) that contribute to a bound sequence’s positive label, similar to key phrase extraction in natural language processing (right).

Our overarching goal is to discover syntactically and semantically meaningful *k*- mers in the context of RBP binding. Our conceptual formulation and structural design choices are based on the emulation of linguistic forms to model RNA sequences with properties and structures analogous to those present in natural languages. With our linguistic representation of RBP binding sites, we modeled genomics as a natural language, treating sentences as sequences, key phrases as significant regions, and words as *k*-mers.

Lexically, we define *k*-mers as words, or the basic units of our RNA lexeme. We consider *k*-mers to have individual roles as syntactic units, and we use the significance or insignificance of *k*-mers to model the relative importance of words within a sentence. Further, we relate the lexical semantics of *k*-mers to their degree of overrepresentation within a dataset, and to their membership as either a component of a motif or an element of the context facilitating binding events.

Syntactically, we define a syntactic form that differentiates context *k*-mers from motif instances and captures the influence of the left and right flanking regions. This syntactic form, which we collectively term the “context”, is composed of three parts: a central *k*-mer (i.e., target) serving the role of a candidate motif instance, and two flanking segments, each consisting of a predetermined number of *k*-mers, serving the roles of the left and right contexts, respectively (**Figure 1B**). We liken the process of differentiating significant regions from insignificant regions to the role of syntactic parsing, and we consider significant regions to be those with important contributions to the sentence’s overall meaning.

### 2.4 A natural language-based sequence decomposition method to represent RBP motifs and contexts

We implemented a novel double sliding window-based method to deconstruct sequences into *k*-mers representing lexical units, then used the lexical units to construct contexts according to our defined syntactic form (**Figure 1B, 1C, 1D**). Let *X* denote the set of sequences belonging to a given RBP, where *X* = {(*X*_1_, *Y*_1_), (*X*_2_, *Y*_2_), …, (*X*_*n*_, *Y*_*n*_)}, such that *X*_*i*_ [1 ≤ *i* ≤ *n*] denotes a sequence and *Y*_*i*_ ∈ *Y* = { 0, 1 } denotes the associated ground-truth class label of *X*_*i*_, with 0 representing the unbound state and 1 representing the bound state.

In the first stage of decomposition (**Algorithm 1**, line 5-6), we apply a sliding window *w*_1_ of size *k*_*target*_ (i.e., the desired length of each target *k*-mer) and stride *s*_*target*_ (i.e., the desired step size of sliding window *w*_1_) over each sequence *X*_*i*_ to deconstruct the sequence into a series of *k*-mers; *w*_1_ yields 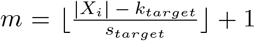 *k*-mers per sequence. In the second stage of decomposition (**Algorithm 1**, lines 7-10), we anchor on a given *k*-mer from the first stage – now designated a target, or candidate RBP binding site – and use a second sliding window *w*_2_ of size *k*_*contexts*_ (i.e., the desired length of the context *k*-mers), stride *s*_*contexts*_ (i.e., the desired step size of sliding window *w*_2_), and span *n*_*contexts*_ (i.e., the desired number of *k*-mers in each flanking region) to construct the left and right context elements for that target. A set composed of a left context, target, and right context is collectively termed a “context” (**Figure 1B**).

We used *k*_*target*_ = 5 due to its demonstration as a robust and representative RBP motif length [3] and *s*_*target*_ = 1 to consider every *k*-mer in a sequence as a potential binding site. We similarly used *k*_*contexts*_ = 5 for context *k*-mers but used *s*_*contexts*_ = 5 to ensure no overlap between successive *k*-mers in the left and right contexts. We used *n*_*contexts*_ = 5 to allow inspection of respectably-sized flanking regions for the analysis of contextual effects. Given our initial sequence length, we have 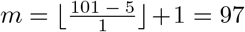 target *k*-mers per sequence, which also corresponds to the total number of contexts per sequence

The set of contexts derived from sequence *X*_*i*_ are denoted *C*_*i*_ = {*C*_*i*1_, *C*_*i*2_, …, *C*_*im*_}, where *C*_*ij*_ represents the context at position *j* [1 ≤ *j* ≤ *m*] of the *i*^*th*^ sequence. We repeat this two-stage sequence decomposition procedure for each sequence *X*_*i*_ in *X*. We denote the full set of contexts for all sequences in *X* as *C* = {(*C*_1_, *Z*_1_), …, (*C*_*n*_, *Z*_*n*_)}, where each *Z*_*i*_ = {*Z*_*i*1_, *Z*_*i*2_, …, *Z*_*im*_} denotes the true labels of the corresponding contexts in *C*_*i*_, which are unknown *a priori* [7].

### 2.5 Formulation of the RBP binding prediction task as a Multiple Instance Learning (MIL) problem

Our approach to sequence decomposition organically led to an MIL-based model in which we frame the RBP binding prediction task as an MIL problem by considering each sequence *X*_*i*_ [1 ≤ *i* ≤ *n*] as a bag and its constituent contexts *C*_*ij*_ [1 ≤ *j* ≤ *m*] as the bag’s instances. Each context *C*_*ij*_ has a corresponding true instance label *Z*_*ij*_ that is unknown *a priori*. The formulated MIL problem is to determine the identity of key instances (i.e., contexts) that significantly contribute to the label of the parent bag (i.e., sequence). We used label inheritance to initialize instance labels such that each context “inherits” the label of its parent sequence (**Figure 1D**, left; **Algorithm 1**, line 11). We sought to predict labels 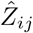 and to use these labels to determine which context(s) are key instances for their corresponding sequences.

We consider two established strategies for bag label assignment (**Supplemental Note .2**). In the standard MIL assumption, a bag is assigned a positive label if it contains at least one positive instance and a negative label if and only if all of its instances are negative [6, 7]. Conversely, the collective MIL assumption considers each bag to be a random representative sample of instances drawn from the bag’s underlying probability distribution; the bag-level class probability is computed using an aggregation-based method to combine instance-level class probabilities, which contribute independently and equally to the bag’s label [6, 7]. We adopted the collective assumption due to the high likelihood that a group of contexts contributes to the bound sequence label.

### 2.6 Solving the MIL problem

The problem at hand is two-fold: (1) accurate determination of instance-level labels, and (2) removal of labeling noise introduced by label inheritance, which could lead to the inclusion of weakly significant contexts. To identify key regions contributing to binding events and to counter the inherent label ambiguity, we required a method that would retain only contexts with strong salient features and remove contexts with weak to no salient features.

To filter out weak contexts, we devised a two-phase strategy termed “iterative relabeling” (**Figure 1D**, right). During the preliminary key instance extraction phase, contexts are scored and their features are compared to the features contained in all sequences in the dataset (**Figure 2**, left; **Algorithm 2**, lines 1-2)). Here, our goal is to approximate how well the contexts (analogous to phrases) contribute to the label of the full-length sequence (analogous to sentences), which is a process similar to key phrase extraction methods that seek to identify the phrase within a sentence that best captures the sentence’s general meaning [12].

**Fig. 2.**
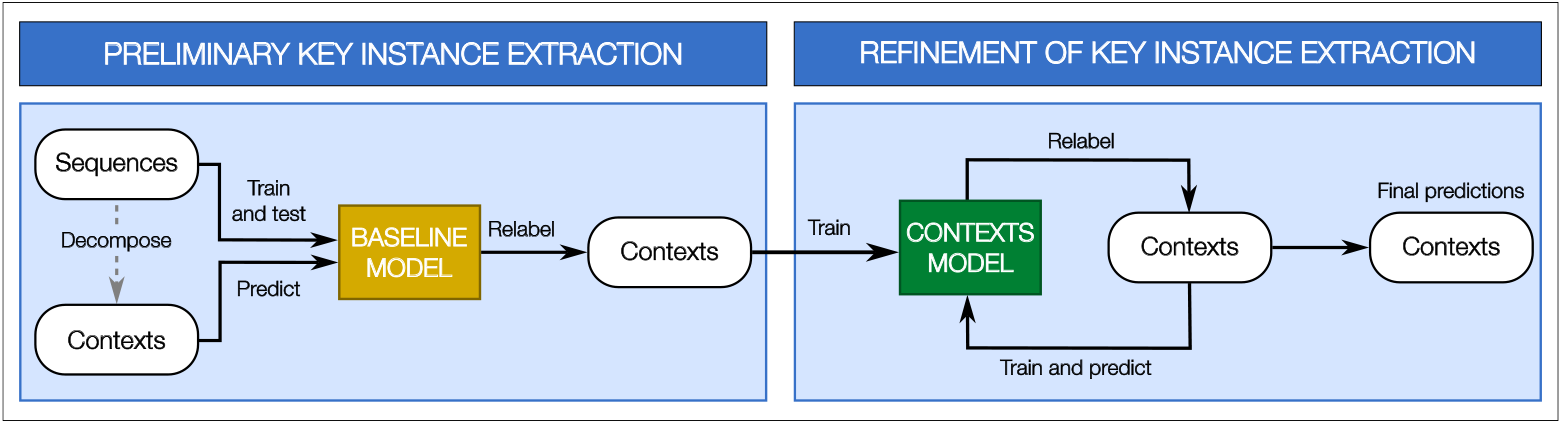
High-level overview of framework workflow. (A) Process diagram of conceptual design. (B) Process diagram of the two-phase key instance extraction process, which corresponds with the two rightmost boxes from panel A.

In the refinement phase, the scored contexts from the preliminary extraction phase are used to fit an initial region (i.e., context) model, after which the context model predictions are used to rescore the regions (**Figure 2**, right; **Algorithm 2**, lines 3-16). The process of fitting the context model and rescoring the contexts is performed iteratively until a satisfactory outcome is achieved (e.g., until convergence, a pre-defined number of iterations *L* is completed, or the first occurrence of overfitting is detected). The refinement phase is similar in principle to reinforcement learning in education settings, in which the learner engages in multiple learning and recall sessions to memorize, discern, and retain the presented content; here, the model learns salient features through reinforcement over multiple iterations, thus allowing the model to discern significant regions with high confidence. This strategy refines the model’s learning of salient features associated with positive observations, thereby ensuring that contexts containing the most salient features are retained while simultaneously sieving out contexts lacking salient features.

### 2.7 Model pre-processing, construction, and training

#### 2.7.1 Data encoding and embedding

We trained a tokenizer on our training dataset to construct a dictionary of all 5-mers, then applied the tokenizer to both the training and test datasets to encode them using either a Bag-of-Words (BoW) or an index-based encoding (indicated below where appropriate). We pre- or post-padded contexts, as needed, to account for those occurring at the beginning or end of a sequence.

#### 2.7.2 Constructing the model architecture

We tested a total of 6 different neural network architectures (**Table 1, Figure S1**). During preliminary testing phases, we conducted experiments with each of these architectures to evaluate differences in prediction performance. We used a hidden layer with the Rectified Linear Unit (ReLu) activation function, a dropout rate of 0.5 to combat overfitting, and the Adam algorithm for optimization [13]. We also used an output layer with a sigmoid activation function as well as the binary cross-entropy loss function, both of which are well-suited for binary classifiers [14].

**Table 1.** Neural network architectures tested.

|  | <u>Abbreviation</u> | <u>Description</u> | <u>Schematic</u> |
| --- | --- | --- | --- |
| 1 | NN-BoW | A shallow neural network comprising a single hidden layer, with a BoW encoding | **Figure S1A** |
| 2 | NN-Embed | A shallow neural network comprising a single hidden layer, with an Embedding layer | **Figure S1B** |
| 3 | CNN | A convolutional neural network (CNN) with max pooling | **Figure S1C** |
| 4 | CNN-LSTM | A CNN with max pooling and long short-term memory (LSTM) | **Figure S1D** |
| 5 | CNN-BiLSTM | A CNN with max pooling and bidirectional LSTM (BiLSTM) | **Figure S1E** |
| 6 | CNN-GRU | A CNN with max pooling and gated recurrent unit (GRU) | **Figure S1F** |

#### 2.7.3 Model training and hyperparameter tuning

Based on our requirements and objectives, our design choices led to two models for each RBP: (1) a reference baseline model, trained and tested on sequences encoded using the BoW method (**Algorithm 3**, lines 2-4), and (2) a context model, trained and tested on contexts encoded using the index-based method (**Algorithm 3**, lines 5-7). For each RBP, we trained binary classification baseline and context models to predict RBP binding at the instance- and bag-levels, using hyperparameter tuning to arrive at the models that yielded the best performance (**Supplemental Note .3**).

#### 2.7.4 Iterative relabeling

We next implemented our iterative relabeling method to filter out weak contexts (**Algorithm 3**, line 8). To implement the preliminary key instance extraction phase, we used the baseline model to score the contexts and obtain initial context predictions, which we used to perform the first relabeling of contexts (**Figure 2**, left; **Algorithm 2**, lines 1-2); this run is designated iteration 0. Because the baseline model was trained and tested on the full-length sequences, we considered the baseline model as our reference and gold standard, serving as a starting measure of comparison between the features within the contexts and those within the full-length sequences.

The refinement phase consists of iterations of context relabeling, in which labels from the previous phase are used to train an initial context model. We then continued updating labels and training context models in an iterative manner by using the predictions from the previous iteration (**Figure 2**, right; **Algorithm 2**, lines 3-16). More specifically, we iteratively score the contexts by using predictions from the model trained at iteration *ℓ* to relabel the contexts for the model’s next training and prediction iteration (*ℓ* + 1). The model at iteration (*ℓ* + 1) is then retrained on the relabeled contexts and used to generate new predictions for the iteration (*ℓ* + 2) of relabeling, and so on. At each iteration, the relabeling policy is applied to contexts originating from positive sequences, such that a context is relabeled 1 if its prediction score is greater than 0.5 and relabeled 0 otherwise. All contexts originating from negative sequences retain their original assigned labels of 0 throughout.

We tested two approaches to train the context models. In the first approach (“refit”), we adopted a single context model for all iterations, gradually retraining this model at each iteration such that its internal parameters are simply updated each time. In the second approach (“new-fit”), we created a new model at each iteration that underwent training using the labeled contexts from the previous iteration, used the new model’s predictions to relabel the contexts again, then discarded the model at the end of the iteration. To optimize runtime, we performed 3 iterations of relabeling.

#### 2.7.5 Evaluation of model performance

Although our datasets were initially balanced, the iterative relabeling process naturally introduced a label imbalance as “weak” contexts from positive sequences were relabeled as negative. We thus used the area under the precision-recall (AUPR) curve to evaluate our prediction accuracy [15]. We computed AUPRs at both the instance- and bag-levels at each stage of training.

We train a contexts model *f*_*C*_ to perform binary classification of the contexts. Formally, bag labels are assigned as follows:

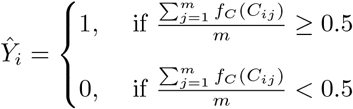

To assess the filtration efficiency of the iterative relabeling process, we computed the witness rates [6] per sequence before and after relabeling. We define the “global witness rate” as the proportion of all contexts in a given RBP’s dataset that were assigned a positive label by the RBP’s model.

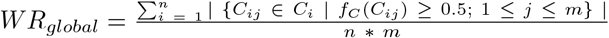

We define the “local witness rate” as the proportion of contexts from a true positive sequence predicted to have a positive label:

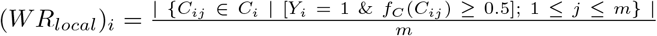

We then performed an average of the local witness rate across all sequences to arrive at a single representative value per RBP:

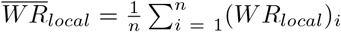

### 2.8 Motif discovery

To validate that our models correctly identify significant regions, we performed motif discovery using STREME from the MEME suite [16]. We first reformatted our contexts predictions in `.fasta` format, then ran STREME using our positively-predicted contexts as our primary set and our negative-predicted contexts as our control set, with the options `--rna --minw 3 --maxw 15 --thresh 0.05 --align center`. To assess the accuracy of motif discovery from our context predictions, we cross-referenced the motifs discovered by STREME with a ground-truth set of 14 RBPs in each cell line whose motifs (1) have been previously established in the literature and (2) are consistent across multiple sources.

## 3 Results

### 3.1 Predictive models exhibit strong performance across RBPs in two cell lines

Our models generally demonstrated a strong performance at each stage of training in both cell lines, with median bag-level AUPRs largely in the mid-to-high eighties (**Table 2, Supplemental Table S2**). We observed that for HepG2, the median final instance- and bag-level AUPRs were 84.5% and 86.5%, respectively (**Table 2A**), and for K562, the median final instance- and bag-level AUPRs were 82.5% and 86.0%, respectively (**Table 2B**). Overall, performance at the instance-level was consistently lower than at the bag-level at each training stage across RBPs (**Figure 3A, 3**B; **Supplemental Table S2**), an expected outcome for a number of reasons. First, ground-truth labels at the instance-level are unavailable, meaning that instance-level performance is not computed relative to a ground-truth. Second, instance-level performance is computed by comparing labels at successive relabeling iterations. Discordance in context labels between successive iterations is expected and intrinsic to the relabeling process, which naturally affects performance but does not necessarily signify an issue, and hence is not an unwanted outcome. Third, fluctuations in instance-level performance between successive iterations are expected as long as there is a noticeable convergence of performance, within an acceptable threshold of overfitting, in the final outcome. We observe that our performance at the instance-level is largely consistent or increasing across relabeling iterations, which is a strong indication of the stability of the relabeling process. Accordingly, we observed that the gap between instance- and bag-level AUPR is reduced substantially after relabeling; in HepG2, the median instance-bag gap is 8.3% before relabeling but only 2% after relabeling, and in K562, the median instance-bag gap is 7.7% before relabeling but only 3.5% after relabeling.

**Table 2.** Statistics of model test performance, in AUPR, at each stage of training in both cell lines. Comprehensive performance metrics from which these statistics were derived are available in **Supplemental Table S2**.

| **(A) HepG2** |  |  |  |  |  |  |
| --- | --- | --- | --- | --- | --- | --- |
| <u>Model</u> | <u>Level</u> | <u>Minimum</u> | <u>Maximum</u> | <u>Median</u> | <u>Mean</u> | <u>Std. Dev.</u> |
| Baseline | Sequence | 0.570 | 0.975 | 0.846 | 0.822 | 0.116 |
| Contexts | Instance | 0.587 | 0.940 | 0.803 | 0.785 | 0.102 |
|  | Bag | 0.633 | 0.983 | 0.886 | 0.856 | 0.0958 |
| Baseline on contexts | Instance | 0.529 | 0.927 | 0.739 | 0.734 | 0.118 |
|  | Bag | 0.578 | 0.980 | 0.852 | 0.825 | 0.115 |
| Contexts, relabeled | Instance | 0.463 | 0.990 | 0.845 | 0.806 | 0.133 |
|  | Bag | 0.596 | 0.981 | 0.865 | 0.841 | 0.105 |
| **(B) K562** |  |  |  |  |  |  |
| <u>Model</u> | <u>Level</u> | <u>Minimum</u> | <u>Maximum</u> | <u>Median</u> | <u>Mean</u> | <u>Std. Dev.</u> |
| Baseline | Sequence | 0.554 | 0.983 | 0.831 | 0.819 | 0.105 |
| Contexts | Instance | 0.588 | 0.938 | 0.796 | 0.781 | 0.0856 |
|  | Bag | 0.641 | 0.990 | 0.873 | 0.858 | 0.0815 |
| Baseline on contexts | Instance | 0.515 | 0.909 | 0.712 | 0.723 | 0.103 |
|  | Bag | 0.525 | 0.982 | 0.835 | 0.820 | 0.107 |
| Contexts, relabeled | Instance | 0.558 | 0.985 | 0.825 | 0.813 | 0.0993 |
|  | Bag | 0.608 | 0.986 | 0.860 | 0.845 | 0.0880 |

**Fig. 3.**
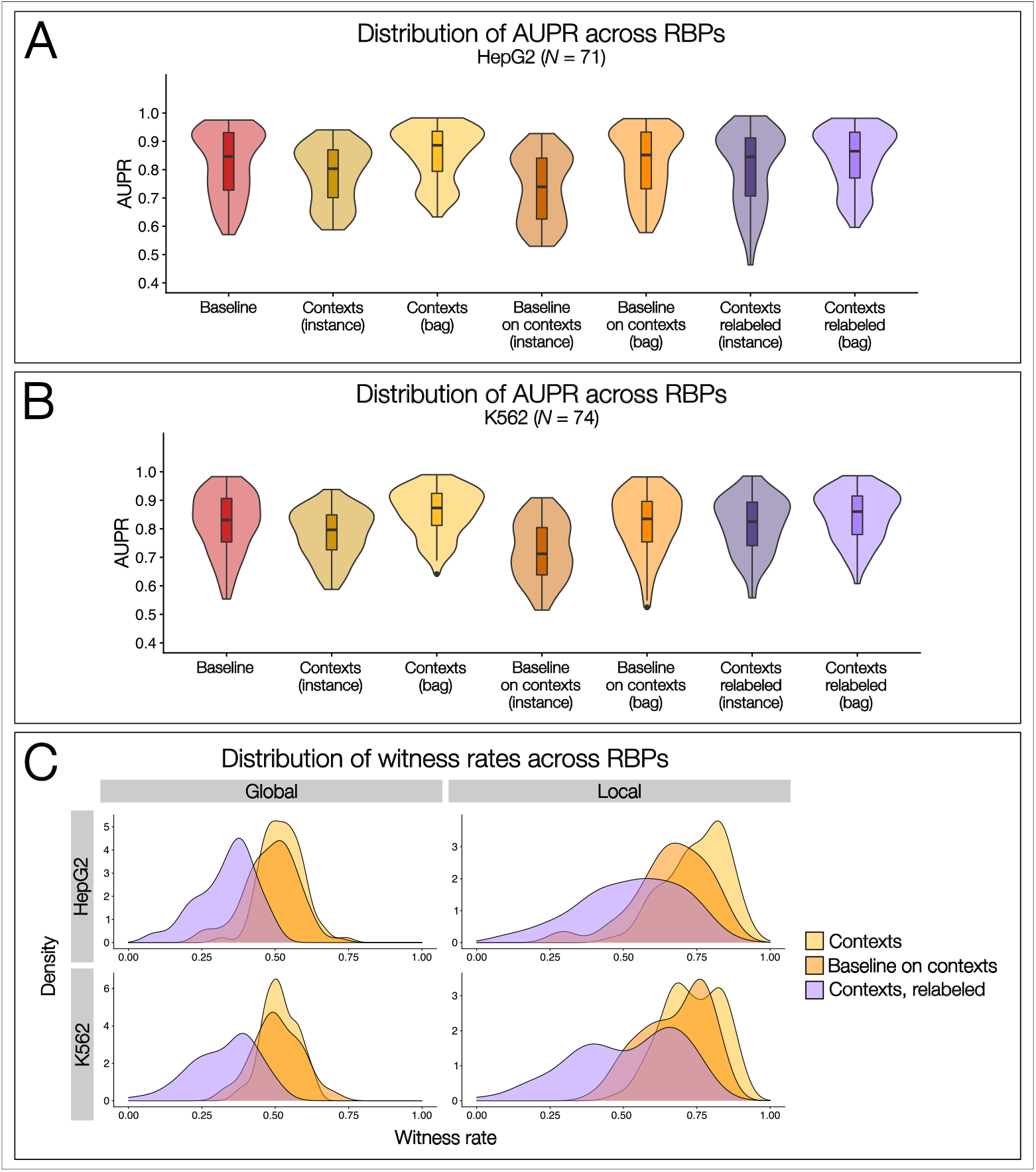
Model performance across all RBPs in both cell lines. Distribution of AUPR across (A) 71 RBPs in HepG2 and (B) 74 RBPs in K562. From left to right: Baseline model’s performance on sequences deconstructed into *k*-mers; Instance- (dark yellow) and bag- (light yellow) level performance of the contexts model before iterative relabeling; Instance- (dark orange) and bag- (light orange) level performance of the baseline model run on contexts; Instance- (dark purple) and bag- (light purple) level performance of the contexts model after iterative relabeling, for the best iteration per RBP. (C) Global (left) and local (right) witness rate distributions at each training stage in HepG2 (top) and K562 (bottom).

The stabilization of the instance-bag gap occurs due to the reinforcement of consistent context labels at each iteration, and it is a further sign that the model’s training had stabilized and neared convergence at the completion of the relabeling process. Therefore, these observations of instance-level performance do not signify that the context model is performing suboptimally or inefficiently, as it would otherwise have negatively affected the accuracy of their respective bag counterparts, which we do not observe. Instead, we consider the instance-level performance not as a measure of true performance of the context models, but instead as a measure of relative performance between two consecutive iterations of relabeling and as a measure of the stability, consistency, validity and efficacy of the relabeling process.

In contrast, we consider bag-level performance as the true measure of context model accuracy because they are computed relative to known ground-truth sequence labels. The bag-level performance was largely stable or increasing across iterations, indicating the stability and efficacy of our predictions during iterative relabeling. Further, together with the instance-level performance, the high bag-level performance indicates that our models achieve good accuracy after iterative relabeling, which serves as additional evidence of the successful removal of weak contexts. Importantly, the distribution of bag-level AUPRs is predominantly preserved across the entire training process (**Figure 3A, 3B**; light yellow, light orange, and light purple violins). Together with the observed reduction in witness rates after iterative relabeling (see next section), the consistency in bag-level performance demonstrates that the contexts model filtered out labeling noise while simultaneously attaining a stable performance.

Finally, bag-level performance of the contexts model at each iteration rivaled or even exceeded that of the baseline model across all RBPs in both cell lines (**Supplemental Table S2**); for example, in HepG2, the median baseline AUPR was 84.6% while the median final bag-level AUPR was 86.5%, and in K562, the median baseline AUPR was 83.1% while the median final bag-level AUPR was 86.0% (**Table 2**). This robust performance indicates that the contexts model was able to learn the same or similar salient features as the baseline model while still filtering noisy contexts (see next section), indicating the efficiency of both the contexts model and iterative relabeling. Further, the spread of the bag-level performance distributions was even smaller than that of the baseline model performance distribution (e.g., in HepG2, baseline and final bag-level standard deviations are 0.116 and 0.105, respectively, and in K562, baseline and final bag-level standard deviations are 0.105 and 0.0880, respectively; **Figure 3A, 3B**, red vs. light purple violin-boxplots; **Table 2**).

In summary, we find that our models yield robust and accurate predictions. Our context model demonstrates stable and efficient performance across iterations at both the instance- and bag-levels, demonstrating the stability and accuracy of our method. Further, the gradual decrease in performance gap between instance- and bag-level AUPRs demonstrates that our iterative relabeling process neared convergence. Finally, as we describe in the following section, our high bag-level performance coupled with the decrease in witness rates observed after iterative relabeling indicate that our models are removing labeling noise while maintaining prediction accuracy.

### 3.2 Witness rate statistics reflect the filtering of noisy labels

To assess the efficacy of iterative relabeling in filtering out noisy contexts, we surveyed the empirical global and local witness rates for each RBP across training stages (**Supplemental Table S3**). Because initial datasets have equal numbers of positive and negative observations, the starting global witness rate for every RBP is 0.5 by definition. We observed a slight increase in median global witness rate in the predictions from the first contexts model (HepG2: 0.526, K562: 0.514), followed by a steady decrease in both the baseline on contexts predictions (HepG2: 0.497, K562: 0.503) and final contexts predictions (HepG2: 0.356, K562: 0.351) (**Table 3**). This decrease is also visible in the distributions of global witness rates over all RBPs at each training stage (**Figure 3C**, left column), which display a leftwards shift at each successive training stage. The distribution shifts were less pronounced when comparing the contexts predictions (yellow) to the baseline on contexts predictions (orange), but more pronounced when comparing the baseline on contexts predictions (orange) to the final contexts predictions (purple). On average, approximately 1*/*3 of all contexts were predicted as positive instances after the final iteration of relabeling (**Table 3**; HepG2: 0.336, K562: 0.328), thereby demonstrating a sizable reduction in global witness rate and, consequently, a sizable degree of context filtering.

**Table 3.**
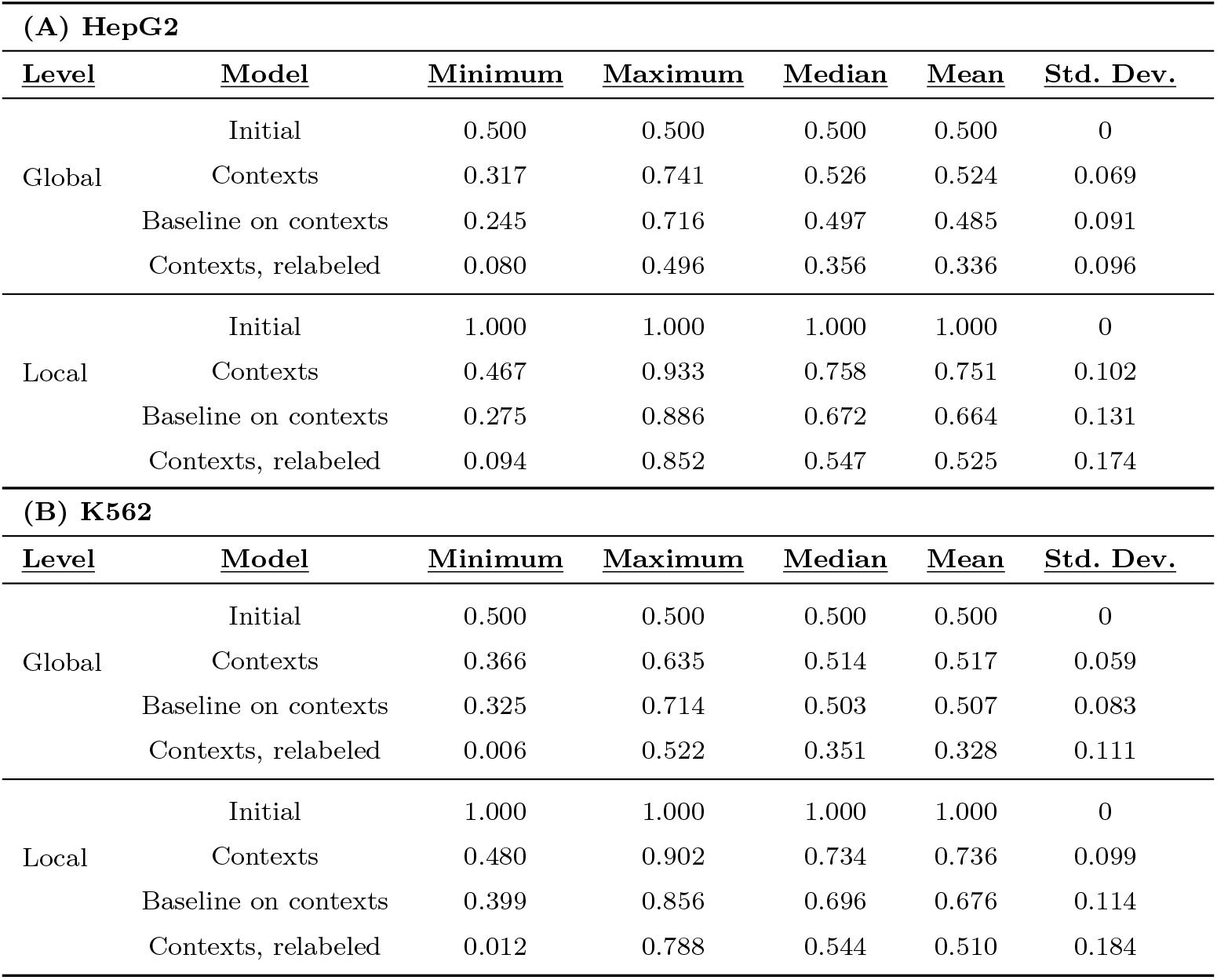
Witness rate statistics across all RBPs. Global and local witness rate statistics at each training stage in (A) HepG2 and (B) K562. Comprehensive witness rates from which these statistics were derived are available in **Supplemental Table S3**.

| **(A) HepG2** |  |  |  |  |  |  |
| --- | --- | --- | --- | --- | --- | --- |
| **<u>Level</u>** | **<u>Model</u>** | **<u>Minimum</u>** | **<u>Maximum</u>** | **<u>Median</u>** | **<u>Mean</u>** | **<u>Std. Dev.</u>** |
| Global | Initial | 0.500 | 0.500 | 0.500 | 0.500 | 0 |
|  | Contexts | 0.317 | 0.741 | 0.526 | 0.524 | 0.069 |
|  | Baseline on contexts | 0.245 | 0.716 | 0.497 | 0.485 | 0.091 |
|  | Contexts, relabeled | 0.080 | 0.496 | 0.356 | 0.336 | 0.096 |
| Local | Initial | 1.000 | 1.000 | 1.000 | 1.000 | 0 |
|  | Contexts | 0.467 | 0.933 | 0.758 | 0.751 | 0.102 |
|  | Baseline on contexts | 0.275 | 0.886 | 0.672 | 0.664 | 0.131 |
|  | Contexts, relabeled | 0.094 | 0.852 | 0.547 | 0.525 | 0.174 |
| **(B) K562** |  |  |  |  |  |  |
| **<u>Level</u>** | **<u>Model</u>** | **<u>Minimum</u>** | **<u>Maximum</u>** | **<u>Median</u>** | **<u>Mean</u>** | **<u>Std. Dev.</u>** |
| Global | Initial | 0.500 | 0.500 | 0.500 | 0.500 | 0 |
|  | Contexts | 0.366 | 0.635 | 0.514 | 0.517 | 0.059 |
|  | Baseline on contexts | 0.325 | 0.714 | 0.503 | 0.507 | 0.083 |
|  | Contexts, relabeled | 0.006 | 0.522 | 0.351 | 0.328 | 0.111 |
| Local | Initial | 1.000 | 1.000 | 1.000 | 1.000 | 0 |
|  | Contexts | 0.480 | 0.902 | 0.734 | 0.736 | 0.099 |
|  | Baseline on contexts | 0.399 | 0.856 | 0.696 | 0.676 | 0.114 |
|  | Contexts, relabeled | 0.012 | 0.788 | 0.544 | 0.510 | 0.184 |

Because label inheritance assigns a positive label to every context originating from a positive sequence, the starting local witness rate is 1.0 by definition. We observed a consistent and substantial decrease in local witness rate at each stage, with median values of 0.758, 0.672, and 0.547 in HepG2 and 0.734, 0.696, and 0.544 in K562 for the initial contexts predictions, baseline on contexts predictions, and final contexts predictions, respectively (**Table 3**). The successive distributional shifts from right to left are also very apparent, demonstrating a clear reduction in positively-labeled contexts after the final iteration of relabeling (**Figure 3**, right column). On average, about 1*/*2 of the contexts from true positive sequences retained their inherited positive label (**Table 3**; HepG2: 0.525, K562: 0.510), with the remainder being filtered out, thus indicating a substantial degree of noisy context reduction.

Finally, the bag performance and witness rate results demonstrate the accuracy and robustness of our models. After relabeling, the bag performance was largely preserved, even as witness rates substantially decreased. Comparing the initial and final median witness rates (**Table 3**), we observe 1.4-fold and 1.8-fold decreases in global and local witness rates, respectively, in both cell lines, indicating that our trained models successfully achieved the removal of labeling noise from positive sequences while simultaneously maintaining bag-level prediction performance.

### 3.3 Standard vs. Collective MIL Assumption

We compared bag performance using both the standard and collective MIL assumptions (**Supplemental Table S4**). The final bag-level AUPRs computed according to the collective assumption were predominantly greater than those computed according to the standard assumption across all RBPs in both cell lines (**Supplemental Table S2, 4**). The median standard bag AUPRs in HepG2 for the initial context predictions, baseline on context predictions, and final context predictions were 0.838, 0.826, and 0.827, respectively (**Table 4A**), all of which are less than the corresponding values for the median collective bag AUPRs (**Table 2A**; 0.886, 0.852, and 0.865, respectively). Similarly, in K562, the median standard bag AUPRs for the three model types were 0.832, 0.805, and 0.824, respectively (**Table 4B**), which were all less than the corresponding median collective bag performance values (**Table 2B**; 0.873, 0.835, and 0.860, respectively).

**Table 4.** Bag performance computed under the standard MIL assumption in (A) HepG2 and (B) K562. To compare and contrast bag performance values with those computed under the collective MIL assumption, refer to **Table 2**. Comprehensive performance metrics computed according to the standard assumption are available in **Supplemental Table S4**.

| **(A) HepG2** |  |  |  |  |  |
| --- | --- | --- | --- | --- | --- |
| <u>Model</u> | <u>Minimum</u> | <u>Maximum</u> | <u>Median</u> | <u>Mean</u> | <u>Std. Dev.</u> |
| Contexts | 0.628 | 0.960 | 0.838 | 0.822 | 0.099 |
| Baseline on contexts | 0.546 | 0.967 | 0.826 | 0.794 | 0.124 |
| Contexts, relabeled | 0.572 | 0.958 | 0.827 | 0.812 | 0.105 |
| **(B) K562** |  |  |  |  |  |
| <u>Model</u> | <u>Minimum</u> | <u>Maximum</u> | <u>Median</u> | <u>Mean</u> | <u>Std. Dev.</u> |
| Contexts | 0.618 | 0.965 | 0.832 | 0.828 | 0.084 |
| Baseline on contexts | 0.519 | 0.979 | 0.805 | 0.790 | 0.109 |
| Contexts, relabeled | 0.603 | 0.962 | 0.824 | 0.821 | 0.085 |

The general distribution of AUPR values for the standard MIL assumption was also consistently shifted downwards compared to the distribution for the collective MIL assumption (**Figure 4A, 4B**). Our findings are corroborated by previous studies, which have similarly demonstrated that the standard assumption often yields poor performance due to noise introduced by label inheritance [6, 7]. The collective assumption, by contrast, often yields better performance due to its equal treatment of instances and because it is less susceptible to labeling noise, particularly for bags belonging to the negative class that contain a few positive instances [6]. These findings support our decision to use the collective MIL assumption.

**Fig. 4.**
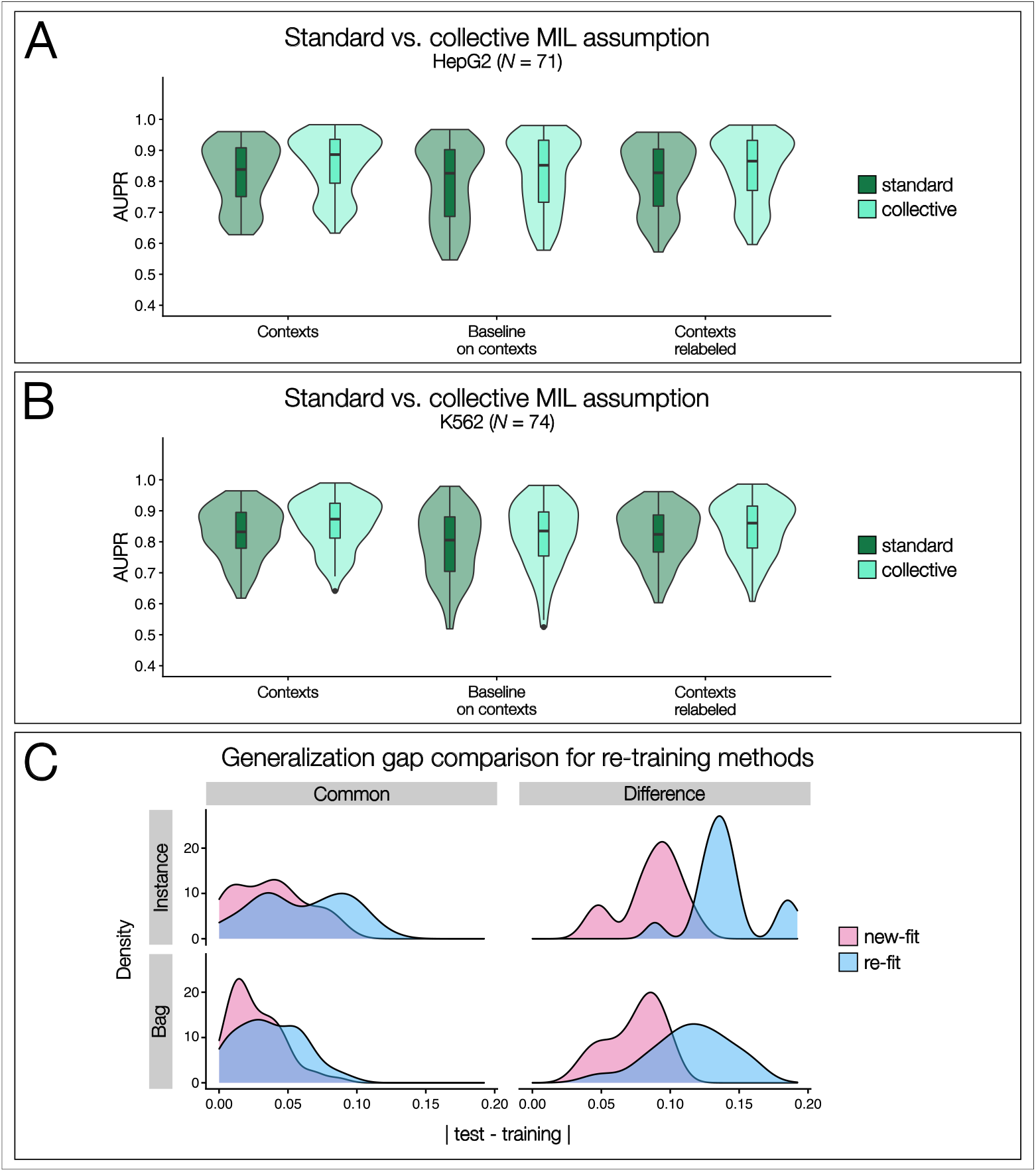
Comparison of different evaluation metrics during model construction and testing. Comparison of bag aggregation methods in (A) HepG2 and (B) K562; distributions of AUPRs calculated in accordance with the standard MIL assumption (dark green) are consistently lower than those calculated in accordance with the collective MIL assumption (aquamarine). (C) Comparison of generalization gap for model re-training methods during iterative relabeling in HepG2. Distribution of the generalization gaps observed at the instance- (top row) and bag- (bottom row) levels are shown, with the “Common” set (left column) representing the set of non-overfitting RBPs common to both training methods (*N* = 54) and the “Difference” set (right column) representing the set of that overfit by the re-fit method but not by the new-fit method.

### 3.4 Comparison of model retraining methods during iterative relabeling

To compare the two methods of model retraining during iterative relabeling, we conducted a survey of overfitting, finding that the new-fit method did not overfit for 71 RBPs while the re-fit method method did not overfit for 54 RBPs, with the entire latter set contained within the former set (“common”) and 17 RBPs overfitting by the re-fit method but not by the new-fit method (“difference”) (**Table 5, Figure 4C**). These findings demonstrate that the new-fit method resulted in a smaller degree of overfitting and more robustly trained models for a greater number of RBPs than the re-fit method. These findings demonstrate that the new-fit method outperformed the re-fit method, having a smaller degree of overfitting and more robustly trained models for a greater number of RBPs.

**Table 5.** Comparison of generalization gap for model retraining methods during iterative relabeling.

| <u>Set</u> | <u>Level</u> | <u>Training Method</u> | <u>Minimum</u> | <u>Maximum</u> | <u>Median</u> | <u>Average</u> | <u>Standard Deviation</u> |
| --- | --- | --- | --- | --- | --- | --- | --- |
| Intersection | Instance | Re-fit | 8.69E-05 | 0.119 | 0.065 | 0.060 | 0.033 |
|  |  | New-fit | 1.47E-03 | 0.091 | 0.036 | 0.038 | 0.026 |
|  | Bag | Re-fit | 4.37E-04 | 0.095 | 0.034 | 0.037 | 0.024 |
|  |  | New-fit | 9.50E-05 | 0.086 | 0.022 | 0.027 | 0.019 |
| Symmetric Difference | Instance | Re-fit | 0.089 | 0.192 | 0.135 | 0.141 | 0.025 |
|  |  | New-fit | 0.041 | 0.120 | 0.092 | 0.086 | 0.022 |
|  | Bag | Re-fit | 0.049 | 0.160 | 0.114 | 0.115 | 0.029 |
|  |  | New-fit | 0.036 | 0.099 | 0.081 | 0.074 | 0.020 |

We also noticed that the generalization gap was generally smaller for the new-fit model at each iteration, but was generally larger for the re-fit model at each iteration. For example, for the 54 RBPs in common, the median bag generalization gap was 0.022 for the new-fit method but 0.034 for the re-fit method; conversely, for the 17 RBPs in difference, the median bag generalization gap was 0.081 for the new-fit method but 0.114 for the re-fit method (**Table 5**). The contrast in generalization gap between the training methods is visually apparent in the shift observed in their respective distributions; this shift is already evident for the 54 RBPs in common (**Figure 4C**, left column, gray) but much more pronounced in the 17 RBPs in difference, where the strong rightwards shift of the re-fit distribution (**Figure 4C**, right column, gray) visually explains the overfitting of these RBPs when iterative relabeling was performed by retraining the same contexts model. Conversely, the new-fit distribution (**Figure 4C**, right column, pink) was shifted leftwards, thus helping to explain why these 17 RBPs were recovered by the new-fit method, as they had overall smaller generalization gaps when iterative relabeling was performed by training a new contexts model and thus did not overfit.

### 3.5 Model performance is architecture-independent

We performed extensive preliminary testing of several neural network architectures and observed that larger, more complex architectures did not outperform smaller, simpler architectures. In fact, performance across these architectures was largely comparable, and the neural network with an embedding layer and a single hidden layer was often the highest-scoring architecture, as exemplified by the representative case study of RBFOX2 in HepG2 (**Table 6**). Accordingly, we decided to use a neural network architecture for our work due to its simplicity and clear comparable performance relative to all other tested architectures. Further, this choice complemented our objectives of maintaining an architecture-independent approach.

**Table 6.** Contexts model performance, in AUPR, across architectures. The 6 different architectures tested are as described in **Section 2.7.2**. NN-BoW: neural network with BoW encoding and single hidden layer; NN-Embed: neural network with embedding layer and single hidden layer; CNN: convolutional neural network; CNN-LSTM: CNN with long short-term memory (LSTM); CNN-BiLSTM: CNN with bidirectional LSTM; CNN-GRU: CNN with gated recurrent unit (GRU).

| Architecture | Contexts (instance) | Contexts (bag) |
| --- | --- | --- |
| NN-BoW | 0.876 | 0.931 |
| NN-Embed | 0.877 | 0.936 |
| CNN | 0.870 | 0.933 |
| CNN-LSTM | 0.870 | 0.932 |
| CNN-BiLSTM | 0.869 | 0.931 |
| CNN-GRU | 0.868 | 0.930 |

### 3.6 Motif and context discovery supports accurate identification of salient regions

We leveraged STREME [16] to search for motifs in our contexts, as previously described. Motifs discovered by STREME from our contexts successfully recapitulated the known motifs for 13 of 14 RBPs in HepG2’s ground-truth set; the sole exception in this set for which STREME did not discover the known motif was PABPN1 (**Figure 5**, left column). In K562, STREME discovered motifs that corresponded with the known motifs for all 14 RBPs in the corresponding ground-truth set (**Figure 5**, right column).

**Fig. 5.**
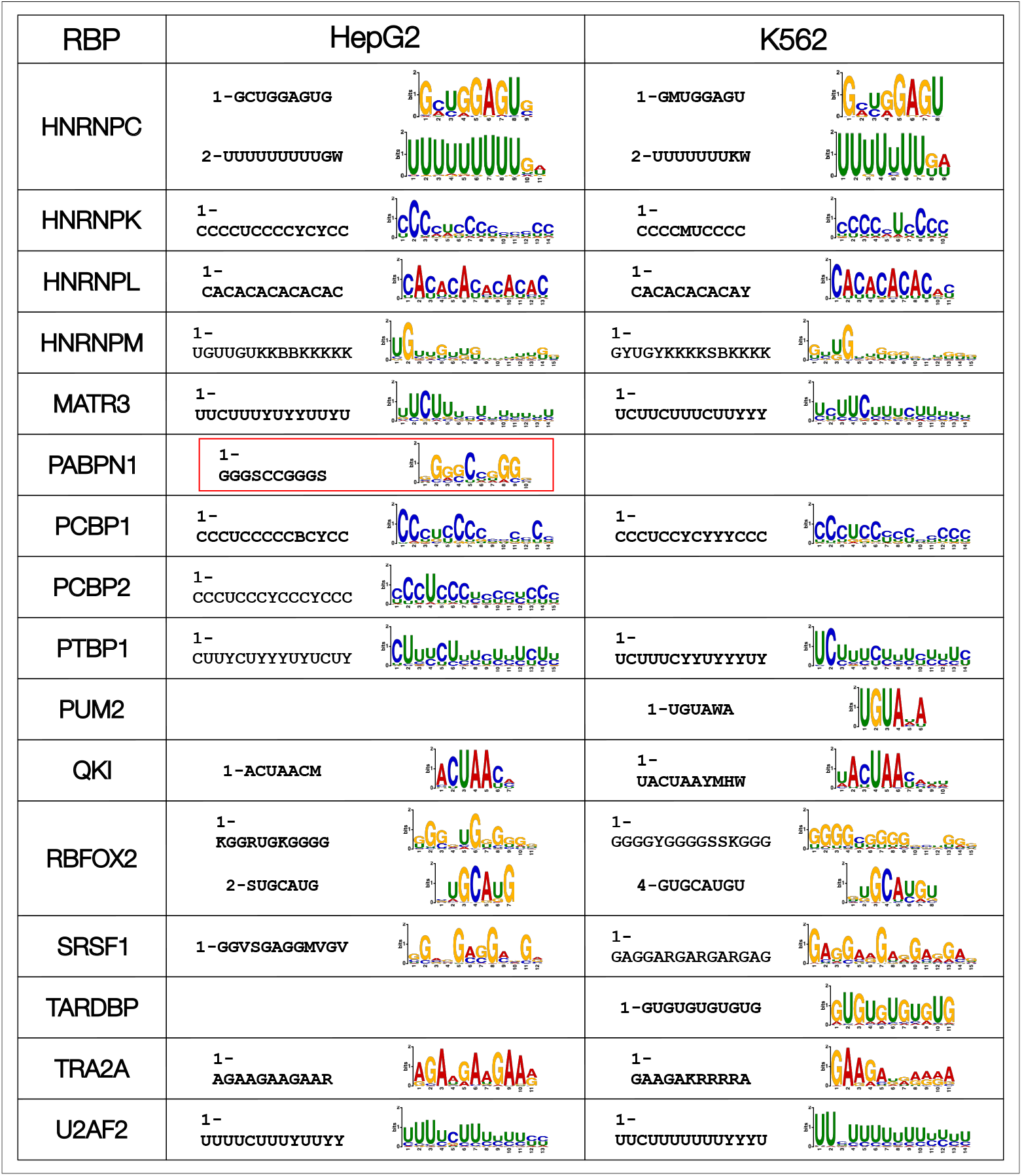
Motif discovery derived from final context predictions for ground-truth sets in HepG2 (left) and K562 (right). STREME was used to discover motifs directly from our final context predictions, returning the top 10 motifs that it discovered. For each RBP, only one motif is shown if the top-ranking STREME motif corresponds with the RBP’s known motif from the literature. If the top-ranking STREME motif does not correspond with the RBP’s known motif, then both the top-ranking and the known motif are displayed (e.g., HNRNPC). PABPN1 in HepG2 (outlined in red) is the only RBP whose STREME motifs do not correspond with the RBP’s known poly(A) motif [21]. Blank cells indicate that eCLIP data was not available for the corresponding RBP in the noted cell line.

Importantly, we also note that STREME did not always return the known motif as its top-ranked motif. For example, the canonical RBFOX2 motif, (U)GCAUG, was ranked 2^*nd*^ in HepG2 and 4^*th*^ in K562, with a G-rich motif being ranked 1^*st*^ in both cell lines; this G-rich motif likely represents the sequence context of RBFOX2, which is known to preferentially bind in G-rich regions [3]. Similarly, the known poly(U) motif for HNRNPC [17] was ranked 2^*nd*^ by STREME, while an alternative GCUGGAGU motif was consistently ranked 1^*st*^. Despite these cases, the STREME-based motif discovery results indicate that our predictive models, and in particular our iterative relabeling strategy, are indeed correctly identifying the key instances (i.e., most highly salient regions), within each sequence that are significant to the binding event and that contribute to the binding motif.

Finally, we also developed a novel consensus-based, deterministic motif and context discovery algorithm [18]. Using the same ground-truth sets as above, we demonstrate that our algorithm recapitulates the known motifs for well-characterized motifs (discovered from our context predictions) with high accuracy, and that our strategy for scoring motifs results in the known motif being ranked as the top hit more often than the results discovered by STREME. Crucially, we used our algorithm to discover RBP binding sequence contexts, and we extensively characterized the flanking nucleotide preferences for all of our RBP datasets in HepG2 and K562 [18]. Taken together, these findings not only demonstrate the validity of our linguistic formulation, conceptualization, implementation, and context predictions presented herein, but it also presents a comprehensive characterization of the *in vivo* RBP binding sequence motifs and contexts for large-scale datasets in two different cell lines, which serves as an important resource for the scientific community in improving our understanding of RBP binding patterns and functions.

## 4 Discussion

In this work, we have presented a novel natural language-based method to model RBP binding prediction comprising (1) a linguistics-inspired representation that introduces lexical, syntactic, and semantic forms to model RNA sequences, (2) a sequence decomposition method that constructs our defined syntactic form to enable the investigation of RBP binding contexts, (3) a weakly supervised Multiple Instance Learning (MIL) formulation of RBP binding which, to the best of our knowledge, is the first application of MIL to the RBP binding prediction task, and (4) a significant region extraction method termed “iterative relabeling” to discover the most highly salient regions significant to RBP binding events, which serves as our solution to the MIL problem. We demonstrate not only that our representation and modeling yield accurate prediction performance, but also that correct binding motifs are discovered from our context predictions, thereby confirming our method’s ability to identify salient regions and their associated motifs. In total, we predicted RBP binding for 71 RBPs in HepG2 and 74 RBPs in K562, and the strong consistency we observe across cell lines at each stage of our work – performance, witness rates, and motif discovery – demonstrate the robustness of our approach.

Our natural language-based representation introduces a number of advantages for modeling RBP binding motifs and contexts. First, the definition of a lexical, syntactic, and semantic structure allows for the clear demarcation of significant regions comprising an RBP binding event: a target, representing a putative RBP binding site, and its context, representing the site’s flanking nucleotides. This representation depicts RBP binding in a biologically relevant manner, aids in distinguishing between direct and contextual effects of RBP binding specificity, and facilitates interpretation. We refrain from making any *a priori* assumptions by allowing each *k*-mer an equal chance of serving as a candidate RBP binding site, which reduces bias, showcases our method’s generalizability, and enables the discovery of secondary motifs that would not otherwise be considered. Our sequence decomposition method is highly customizable and flexible, thus enabling investigations of RBP binding patterns at increased granularity. Finally, a key component of our approach is its high resolution: our context structure allows us to obtain a prediction for every context within a sequence, thus allowing us to pinpoint the exact regions bound by or contributing to an RBP’s binding.

Our iterative relabeling strategy, presented as a machine learning-based implementation, was essential to removing the labeling noise inherent to weakly supervised frameworks like MIL. The gradual filtering of weak contexts, along with the strong model performance presented herein, showcase our approach’s ability to achieve key instance extraction with very good accuracy. It also demonstrates the strength and robustness of our MIL models to filter out label ambiguity introduced by label inheritance. In addition, the syntactic format of the discovered significant regions greatly reduces downstream processing and gives our sequences a defined structure to guide further analyses.

There are several aspects of this work that we believe are worth revisiting in the future. First, extensive investigation of the selection of sliding window parameters will be most informative in determining how *k*-mer and stride sizes effect model performance, significant region extraction, and motif discovery. Similarly, testing of different context lengths would help to reveal the optimal context size necessary for binding, which we hypothesize to be RBP-specific. Bipartite motifs [3] may also be investigated through the introduction of gaps between *k*-mers. We also intend to expand the scope of syntactic analyses within the left and right contexts, which will aid in a better understanding of RBP functions.

Second our approach to hyperparameter tuning balanced a tradeoff between performance and runtime. Hyperparameter tuning is a combinatorially complex NP-hard problem, making exhaustive testing computationally infeasible. We employed a best practice alternative to onlytest randomly selected sets of hyperparameter values, which achieved very good results in a reasonable timeframe (**Supplemental Note .3**). Performance optimization is an architectural aspect of model training that could be further explored.

Third, semi-supervised learning (SSL), a class of weak supervision, would serve as a complement to the iterative relabeling strategy that works from a bottom-up manner, in that the vast majority of observations in the dataset are initially unlabeled but incrementally assigned new labels as model training progresses [19]. We view this as an important area of research that we aim to continue pursuing in the future.

In summary, we were able to develop a robust method that satisfied our project objectives, achieving *high resolution, interpretability*, and *flexibility* by enabling the study and analysis of RBP binding motifs, contexts, and preferences at different levels of granularity through our linguistic-based representation. Further, our method enables *easy access to relative and absolute genomic coordinates*, which facilitates downstream gene- and loci-specific analyses [18], and our method is *architecture-independent*. In conclusion, we present a novel representation and modeling of RBP binding based on linguistics and natural language that robustly predicts RBP binding across cell lines, resulting in accurate RBP motif discovery and enabling the study of RBP binding contexts.

## Conflicts of interest

Zhiping Weng co-founded Rgenta Therapeutics, and she serves as a scientific advisor for the company and is a member of its board.

## Funding

This work was supported by the National Institutes of Health [U24HG012343 to Z.W.].

## Data availability

Data is publicly available at the ENCODE portal [8, 9, 20]. Accessions for eCLIP data used in this study are detailed in **Supplemental Table S1**.

## Code Availability

Code is publicly available at https://github.com/orbitalse/novel RBP discovery methods.

## Author Contributions

S.I.E.: Problem Formulation, Conceptualization, Data Curation, Data Processing, Investigation, Formal Analysis, Methodology, Visualization, Writing of Manuscript (original draft), Review of Manuscript, Editing of Manuscript. Z.W.: Research Area, Discussions, Review of Manuscript. All authors reviewed and approved the final version of the manuscript.

## Supplemental Notes

### 1 On the exponential nature of *k*-mer enumeration

Let *k* denote the maximum word size in a dictionary; the associated vocabulary size is 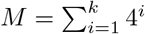. Further, if we let *n* denote the desired sequence length, then we have *M*^*n*^ possible sentences. To demonstrate a simple scenario, let *k* = 5 and *n* = 5. It then follows that the vocabulary size is *M* = 4 + 4^2^ + 4^3^ + 4^4^ + 4^5^ = 1364 words, and the number of possible sentences is subsequently 1364^5^ = 4, 721, 411, 479, 245, 824; one can imagine how quickly this number will grow if both *k* and *n* are increased (**Figure S2**). For example, the vocabulary size is 1, 398, 100 for *k* = 10, and 1, 431, 655, 764 for *k* = 15. The challenge is further exacerbated by the strong possibility that only a small subset of words from the enumeration of each value of *k* are actually syntactically and semantically meaningful within a given biological context.

### 2 Formal definition of the MIL assumptions

Here, we formally define the standard and collective assumptions that are used for bag label assignment. All mathematical notation is consistent with definitions in the main text.

#### Standard MIL assumption

In the standard MIL assumption, a positive label is assigned to a bag if at least one of its constituent instances is positive [22]. Formally, a sequence *X*_*i*_’s bag class label *Y*_*i*_ is defined as:

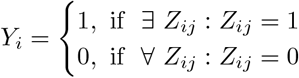

That is, a positive bag has at least one positive instance, but a negative bag comprises only negative instances.

In our formulation, in which true instance labels *Z*_*ij*_ are unknown, instance-level class probabilities predicted by the model *f*_*C*_ are used to reach an approximation of *Z*_*ij*_, which are then used to approximate *Y*_*i*_:

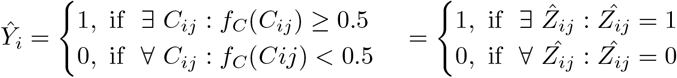

#### Collective MIL assumption

Conversely, the collective MIL assumption regards each constituent instance as an equal contributor to the bag label, and the bag label itself is assigned based on some method of aggregation over the constituent instances [22]. We demonstrate in **Section 3.3** that the standard assumption is outperformed by the collective assumption, an observation that corroborates findings previously reported in the literature [6, 7]. We thus adopted the collective MIL assumption for this work.

We operate under the collective assumption’s postulates [22]; namely:

1. A bag is not a finite set of elements.

2. A bag has an underlying probability distribution from which its constituent instances are randomly sampled.

Let *B* denote a random variable representing a bag, and let *X* denote a random variable representing an instance. Bags and instances are distributed according to some unknown joint probability distribution, *P* (*B, X*), where:

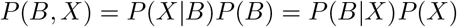

Solving for *P* (*X*|*B*), we arrive at:

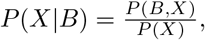

where *P* (*X* |*B*) represents a conditional probability distribution to model a fixed bag over the instance space; i.e., given a bag, what is the probability that a given instance belongs to that bag? Bags can be constructed by sampling from the conditional probability distribution *P* (*X* | *B*); thus instances are assigned to a bag by random sampling from this underlying distribution.

Next, let *Y* denote a random variable representing a binary class label, and let *X* represent, as previously stated, a random variable representing an instance. Instances and class labels are similarly distributed according to some unknown joint probability distribution, *P* (*Y, X*). Labels can thus be assigned to instances based on the unknown conditional distribution *P* (*Y* | *X*).

Now, let *c* be a binary class label such that *c* ∈ { 0, 1 }, let *b* denote a given bag, and let *y* denote the class label of *b*. A given instance *x* from bag *b* is assigned a positive label if *p*(*c* = 1 | *x*) is greater than 0.5, and assigned a negative label otherwise. Under the standard MIL assumption, bags would be assigned labels as follows:

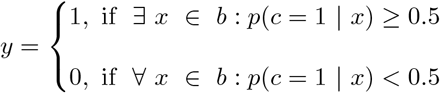

That is, a bag is assigned a label of 1 if there exists at least one instance in that bag with a positive label, and assigned a label of 0 if all instances in that bag have a negative label.

Conversely, under the collective MIL assumption, the probability that a bag *b* has label *c, p*(*c* | *b*), can be approximated by the expected class value of the bag’s population [22].

The general formula for expected value of a continuous random variable is:

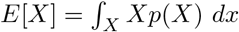

and the conditional expectation is:

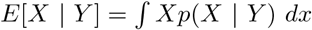

To approximate the bag label by the expected class value of the bag’s population, it follows that:

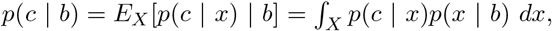

where *x* is an instance and *E*_*X*_ [*p*(*c* | *x*) | *b*] represents the mean of the class probabilities for each instance *x* in a given, fixed bag *b*.

Since *p*(*x* | *b*), the conditional probability distribution of the bag, is unknown, and because we don’t have any way of approximating it, computing *p*(*x* | *b*) is not feasible. A third assumption is added that each instance in a bag contributes equally and independently to the bag’s label by approximating 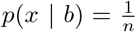; in other words, we approximate *p*(*x* | *b*) by a uniform distribution.

Since *p*(*c* | *x*) can be approximated (in this case, by our MIL model predictions), we instead use the sample of instances in the bag (rather than the bag’s true population distribution) to approximate *p*(*c* | *b*):

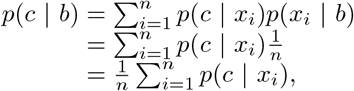

where *n* is the total number of instances in bag *b*. Thus, to approximate bag labels, we average the class probabilities of all its constituent instances.

In summary, the collective MIL assumption has the following key characteristics:

1. **Probabilistic**: rather than being viewed as a fixed collection, a bag is instead considered to be a random representative sample of instances drawn from the bag’s underlying probability distribution.
2. **Equal weighting**: each instance contributes independently and equally to the bag’s label.
3. **Aggregation-based**: to determine the final bag label, instance-level class probabilities are combined using some aggregation function; for example, using the average of instance-level class probabilities to compute the bag-level class probability, where a bag is assigned a positive label if the average is greater than 0.5 and a negative label otherwise. This contrasts with the standard assumption, in which bag labels are assigned using the maximum function based on a pre-defined prediction value threshold.
4. **Flexibility**: the collective assumption is a flexible alternative to the standard assumption, as it is easily adapted to problems where the model is threshold-based, or in applications where it is unreasonable to assume that the presence of a single positive instance definitively denotes that the parent bag is positive.

### 3 On hyperparameter tuning

A robust high-performing model is dependent on many aspects of the neural network itself; namely, the size of the neural network (which determines the number of parameters to be trained), the number of hyperparameters associated with the neural network, and the range of values to consider for each hyperparameter. Because hyperparameter tuning is a combinatorially complex problem which only increases in complexity as the number of hyperparameters grows larger, and exhaustive testing of every possible combination of hyperparameter values would be computationally infeasible, we opted for a common alternative: first, devise a method that reduces this computational complexity by instead selecting a range of values for each hyperparameter (after extensive preliminary analyses and experimentation to gauge satisfactory minimum and maximum values for each range); then, apply the method by selecting a combination of values with which to train the model. The combination of hyperparameter values that yields the best-performing model and satisfies overfitting criteria is ultimately selected for the final model. We implemented such an approach by defining the following ranges for each hyperparameter of our network, having arrived at these ranges after considerable experimentation and analyses.

- Embedding dimension: [8, 16, 32, 64, 128]
- Number of neurons: [16, 32, 64, 128, 256, 512, 1024, 2048]
- Batch size: [32, 64, 128, 256, 512, 1024, 2048]
- Number of epochs: [2, 3, 4, 5, 6, 7, 8, 9, 10]

We employed the RandomizedSearchCV utility from scikit-learn [23] to test 25 random combinations of hyperparameter values. To determine which set of hyperparameter values performed optimally for both the baseline and context models, respectively, we defined and applied a set of criteria that balanced a model’s generalization gap (i.e., the difference in training and test performance) with its overall test performance. We first began by identifying any tuning iterations whose generalization gap was less than a predefined threshold of 0.005. If no tuning iterations met this threshold, we then incrementally increased the threshold until at least one tuning interaction satisfied the threshold, which enabled us to isolate the tuning iterations with the smallest generalization gap. Next, from this set of tuning iterations, we designated the one with the maximum test performance as the tuning iteration with the best performance. Finally, we used the optimized hyperparameter values from the selected iteration to train the final model on the full training dataset.

#### Algorithm 1

Sequence Decomposition into *k*-mers and Contexts (*X, w*_1_, *w*_2_)

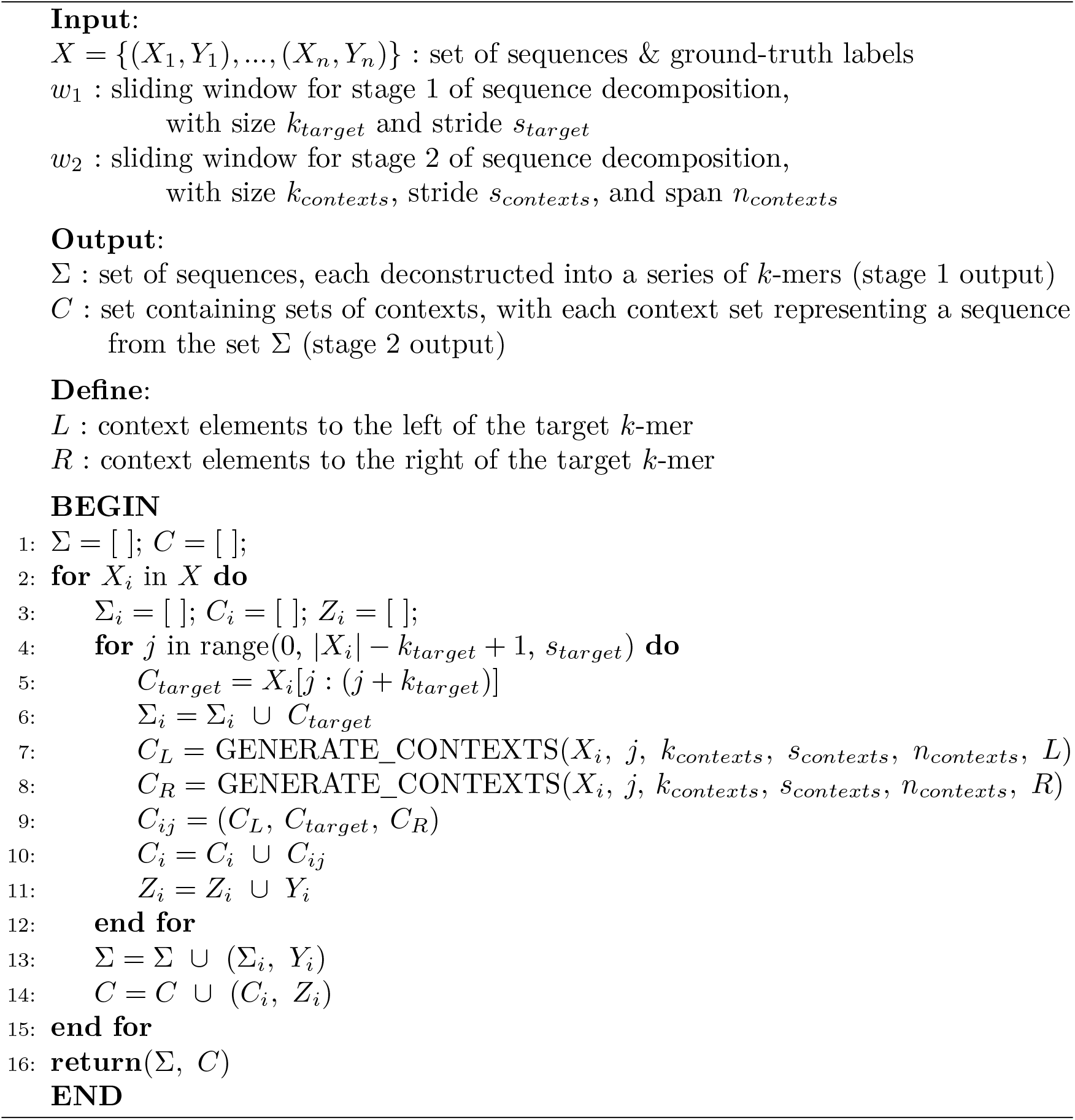

#### Algorithm 2

Iterative Relabeling (*C*, Σ_*V*_, *f*_*B*_, *f*_*C*_, *δ, I, α*)

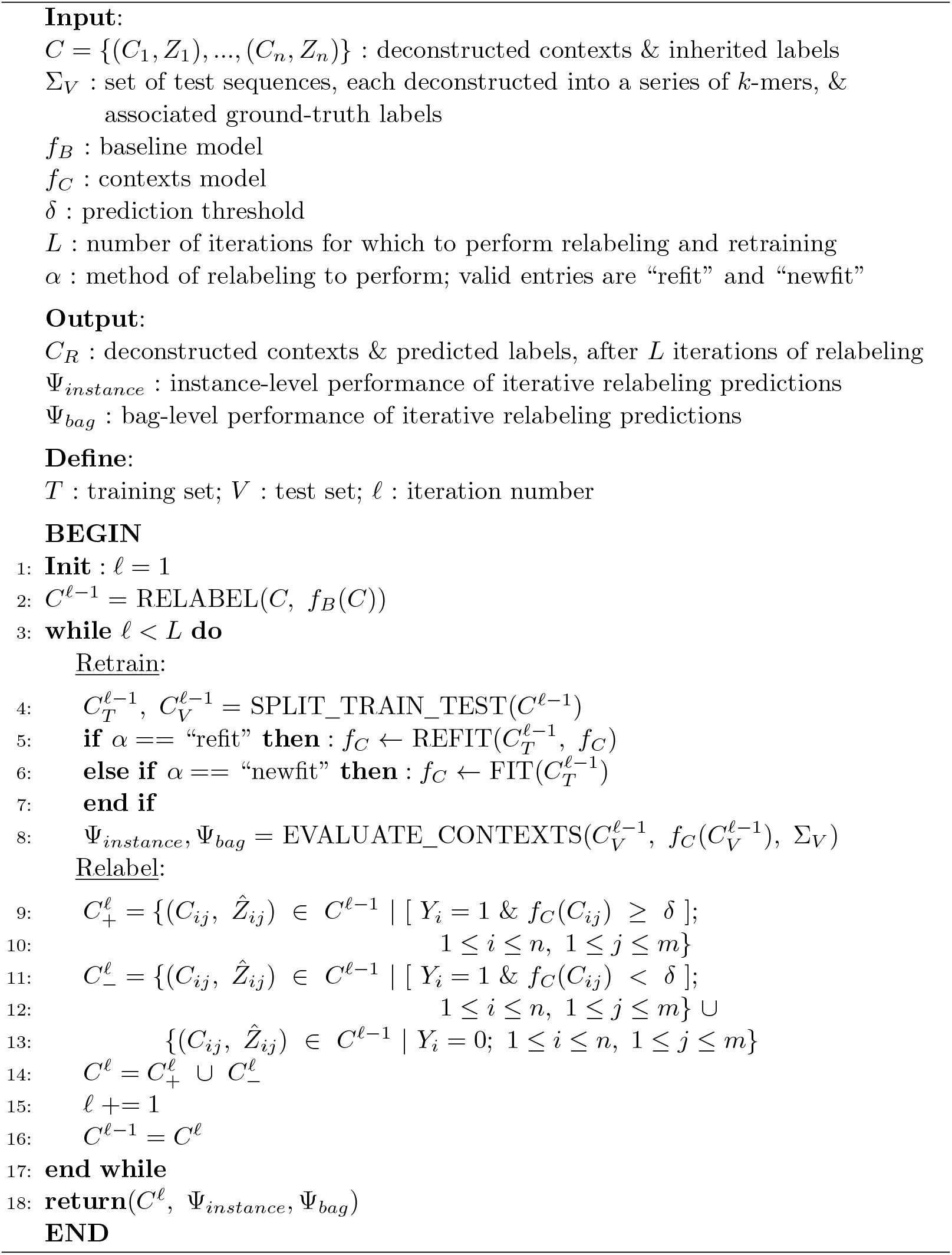

#### Algorithm 3

Multiple Instance Learning Workflow (*X, w*_1_, *w*_2_, *δ, I, α*) (NN-BoW, NN-Embed, CNN, CNN-LSTM, CNN-BiLSTM, CNN-GRU)

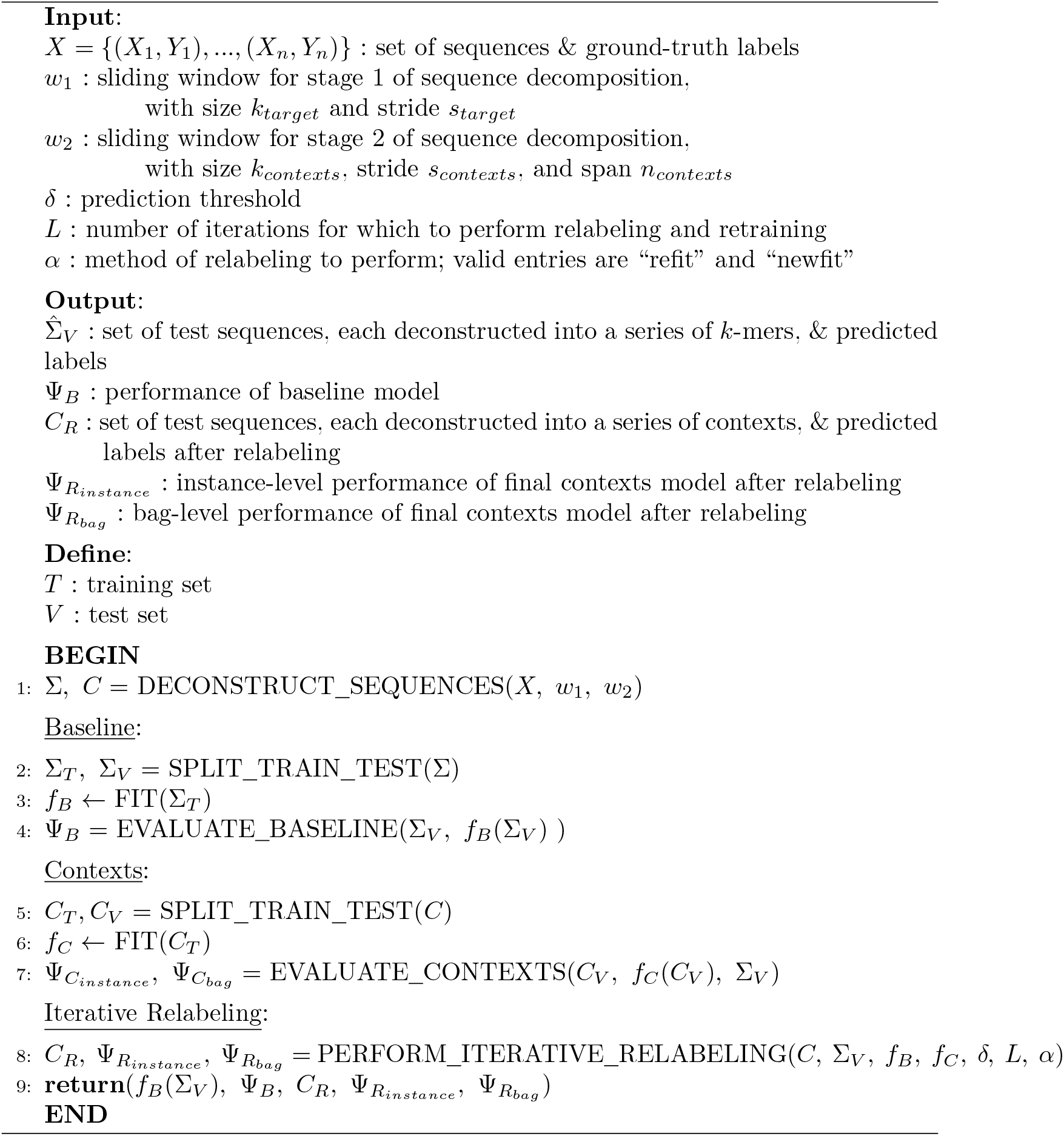

---

**Figure S1.**
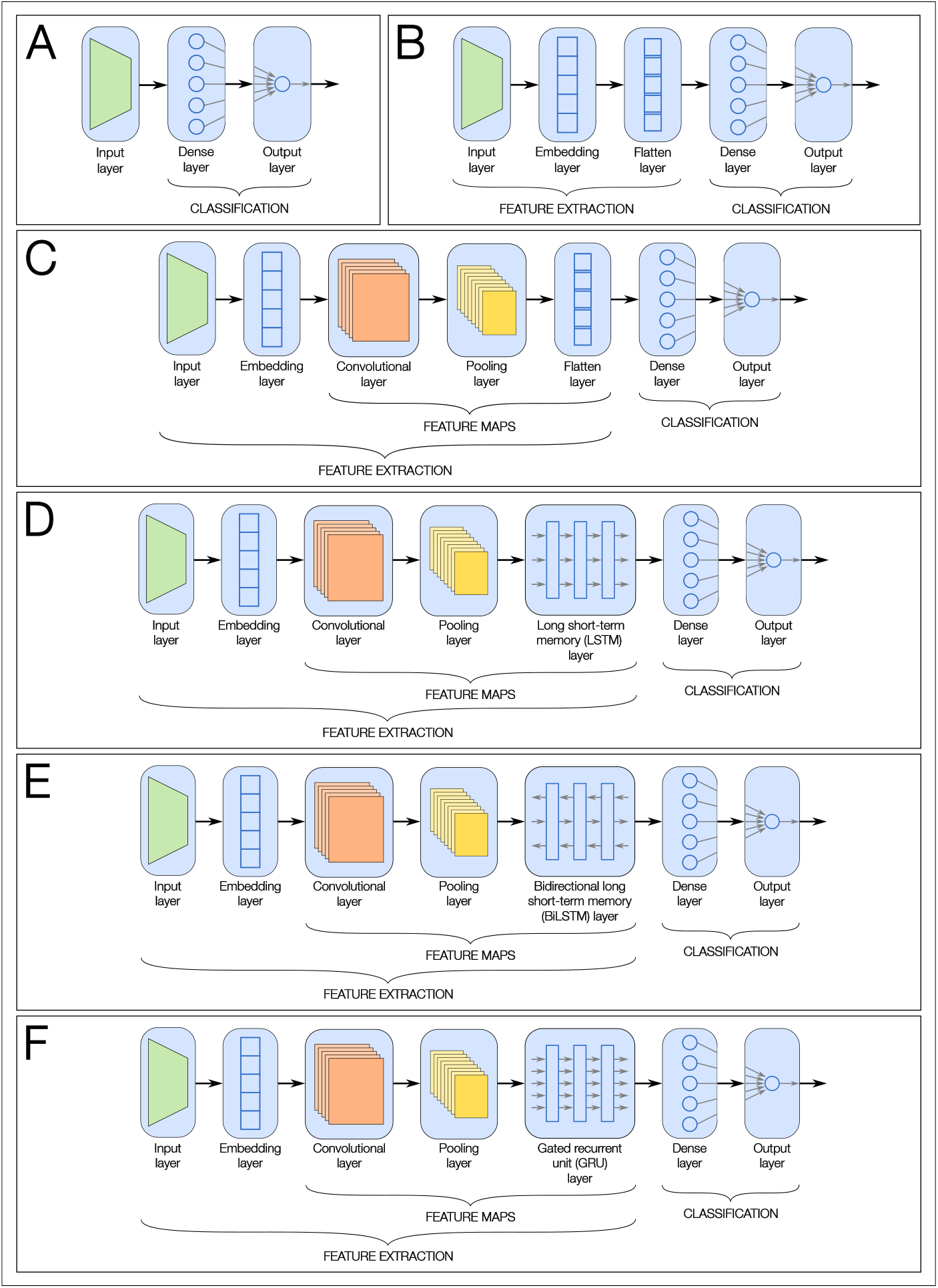
Neural network architectures tested. (A) NN-BoW: a shallow neural network comprising a single hidden layer, with a Bag-of-Words encoding. (B) NN-Embed: a shallow neural network comprising a single hidden layer, with an Embedding layer. (C) CNN: a convolutional neural network (CNN) with max pooling. (D) CNN-LSTM: a CNN with max pooling and long short-term memory (LSTM). (E) a CNN with max pooling and bidirectional LSTM (BiLSTM). (F) CNN-GRU: a CNN with max pooling and gated recurrent unit (GRU).

**Figure S2.**
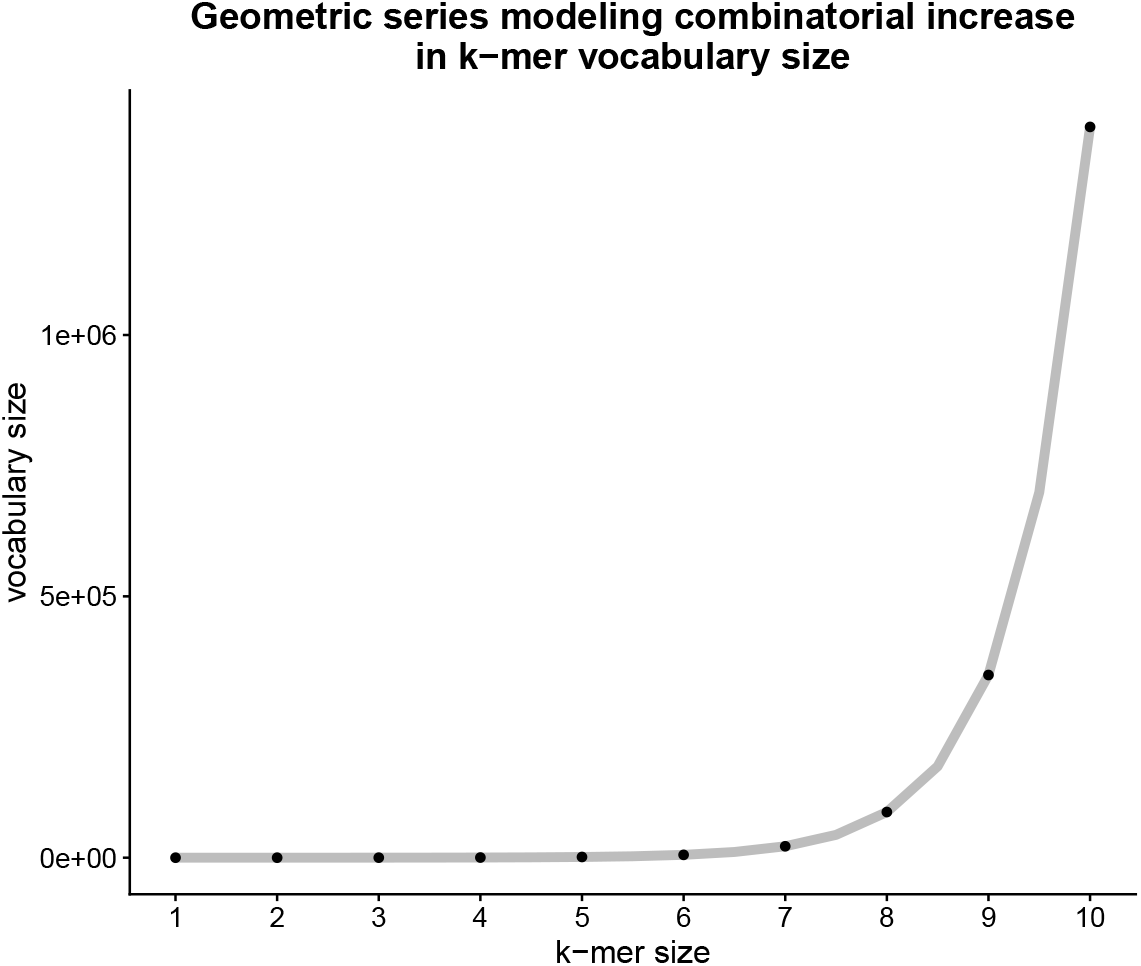
Geometric series modeling increase in *k*-mer vocabulary size. *k*-mer combinatorics can be modeled by the geometric series 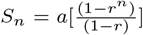, where *a* = 4 and *r* = 4, resulting in the function 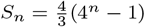.

